# Stress decreases serotonin tone in the nucleus accumbens to promote aversion and potentiate cocaine preference via decreased stimulation of 5-HT_1B_ receptors

**DOI:** 10.1101/2021.06.27.450100

**Authors:** Harrison M Fontaine, Phillip R Silva, Carlie Neiswanger, Rachelle Tran, Antony D Abraham, Benjamin B Land, John F Neumaier, Charles Chavkin

**Affiliations:** Department of Pharmacology, University of Washington School of Medicine, Seattle, WA 98195; Center for the Neurobiology of Addiction, Pain, and Emotion, University of Washington School of Medicine, Seattle, WA 98195; Graduate Program in Neuroscience, University of Washington School of Medicine, Seattle, WA 98195; Department of Psychiatry and Behavioral Sciences, University of Washington School of Medicine, Seattle, WA 98195; Puget Sound VA Health Care System, Seattle, WA 98108

## Abstract

Stress-induced release of dynorphins (Dyn) activates kappa opioid receptors (KOR) in monoaminergic neurons to produce dysphoria and potentiate drug reward; however, the circuit mechanisms responsible for this effect are not known. We found that conditional deletion of KOR from *Slc6a4* (SERT)-expressing neurons blocked stress-induced potentiation of cocaine conditioned place preference (CPP). Within the dorsal raphe nucleus (DRN), two overlapping populations of KOR-expressing neurons: *Slc17a8* (VGluT3) and SERT, were distinguished functionally and anatomically. Optogenetic inhibition of these SERT^+^ neurons potentiated subsequent cocaine CPP, whereas optical inhibition of the VGluT3^+^ neurons blocked subsequent cocaine CPP. SERT^+^/VGluT3^−^ expressing neurons were concentrated in the lateral aspect of the DRN. SERT projections from the DRN were observed in the medial nucleus accumbens (mNAc), but VGluT3 projections were not. Optical inhibition of SERT^+^ neurons produced place aversion, whereas optical stimulation of SERT^+^ terminals in the mNAc attenuated stress-induced increases in forced swim immobility and subsequent cocaine CPP. KOR neurons projecting to mNAc were confined to the lateral aspect of the DRN, and the principal source of dynorphinergic (Pdyn) afferents in the mNAc was from local neurons. Excision of *Pdyn* from the mNAc blocked stress-potentiation of cocaine CPP. Prior studies suggested that stress-induced dynorphin release within the mNAc activates KOR to potentiate cocaine preference by a reduction in 5-HT tone. Consistent with this hypothesis, a transient pharmacological blockade of mNAc 5-HT_1B_ receptors potentiated subsequent cocaine CPP. 5-HT_1B_ is known to be expressed on 5-HT terminals in NAc, and 5-HT_1B_ transcript was also detected in *Pdyn*^+^, *Adora2a*^+^ and *ChAT*^*+*^ (markers for direct pathway, indirect pathway, and cholinergic interneurons, respectively). Following stress exposure, 5-HT_1B_ transcript was selectively elevated in *Pdyn*^*+*^ cells of the mNAc. These findings suggest that Dyn/KOR regulates serotonin activation of 5HT_1B_ receptors within the mNAc and dynamically controls stress response, affect, and drug reward.

## Introduction

Stress has profound effects on the risk of substance use disorders and relapse in humans and promotes drug seeking behaviors in animal models of addiction [1–4]. Animal studies have shown that the endogenous opioid dynorphin (Dyn)^(**footnote 1**)^ and its cognate receptor, the kappa opioid receptor (KOR), are critical to the enhancement of each stage in the progression towards drug addiction, from initial preference, to escalation, and ultimately reinstatement [2,5–7]. These have been shown to be mediated in part by stress-induced modulatory effects on the serotonin (5-HT) system; however, the contribution of serotonin (5-HT) to hedonic processing remains controversial [8–10]. In humans, polymorphisms in genes encoding dynorphin, KOR, and the serotonin transporter (SERT) have been linked to stress-induced depression and increased risk for addiction [11–14].

Stress-evoked release of neuropeptides including corticotropin-releasing factor (CRF) and the prodynorphin-derived peptides impinge on affective circuitry to orchestrate changes in both neurophysiological state and observable behavior [15]. CRF-induced Dyn release is necessary for the dysphoric properties of stress, and Dyn action at KOR on dopaminergic and serotonergic neurons is necessary for a stress-induced dysphoric state, which may underlie stress-potentiation of drug-seeking behaviors [15,16]. KOR activation within serotonergic neurons of the dorsal raphe nucleus (DRN), which is a hedonic hot spot and primary source of forebrain serotonin, results in somatic hyperpolarization and increases the surface expression of SERT in axon terminals projecting to the nucleus accumbens (NAc) [9,10,17,18]. Together, these findings suggest that stress-induced activation of the Dyn-KOR-5-HT axis reduces serotonin tone in the NAc to increase drug reward in mice.

Direct manipulation of serotonergic neuron activity in DRN via optogenetic and chemogenetic techniques, however, has resulted in conflicting conclusions concerning the role of 5-HT in mediating responses to rewarding, aversive, and stressful stimuli [19–24]. These discrepancies may be due to the genetic and anatomical complexity of the DRN as well as the impact of different assay conditions and event timing on stress and reward processing [25,26]. In the present study, we resolved a KOR-expressing, serotonergic projection from the lateral aspect of the DRN to the medial NAc (mNAc) that controls 5-HT tone to regulate stress response, aversion, and reward potentiation. We further implicate presynaptic dynorphin and postsynaptic 5-HT_1B_ receptors within the mNAc in mediating these effects.

## Materials and Methods

### Animals

Adult male C57BL/6 mice and transgenic strains on C57BL/6 background were group housed with access to food and water *ad libitum*. All animal procedures were approved by the University of Washington Institutional Animal Care and Use Committee and conformed to US National Institutes of Health guidelines.

### Behavior

rFSS, cocaine CPP, and social approach were performed as previously described [1,27]*Optical stimulation during rFSS:* SERT^DRN-NAc^ ChR2 mice were tethered throughout swim sessions and stimulated on day 1 and swims two and four of day 2. *DRN inhibition pretreatment*: VGluT3^DRN^ or SERT^DRN^ SwiChR groups received optical inhibition for 30min, followed by cocaine conditioning 30min later. *SERT*^*DRN-NAc*^ *terminal excitation during U50488 pretreatment*: 1hr prior to cocaine conditioning, mice received U50488 and optical stimulation, terminating 5min before cocaine administration. *5-HT*_*1B*_ *antagonist pretreatments:* GR 127935 was infused into the NAc 135min and/or 75min prior to cocaine conditioning or social approach.

### Histology

Immunohistochemistry and RNAscope were performed as previously reported [28,29].

### Data Analysis

The assumption of normal distribution was tested and corrected for when not met. T tests were unpaired, two-two tailed. *Post-hoc* tests were Sidak’s or Dunnett’s where appropriate, with *α*=0.05.

## Detailed methods are provided in Supplemental Materials

### Results

#### KOR expression in SERT neurons is required for stress potentiation of cocaine reward

*Slc6a4-Cre* (‘SERT-Cre’) and *Oprk1-lox/lox* (‘KOR-flx’) mice were crossed as previously described [16], resulting in conditional excision of KOR in SERT^+^ cells (‘KOR^SERT^cKO’) (Figure 1A). These mice and their littermate controls were subjected to repeated forced swim stress (rFSS) prior to cocaine place preference conditioning (CPP) (Figure 1B). Control mice developed significant cocaine CPP, and rFSS induced potentiation of CPP that was absent in KOR^SERT^ cKOs (Figure S1A). To isolate the effect of stress on cocaine preference, we normalized within each genotype to the ‘No rFSS’ preference. Stress increased cocaine preference in controls by more than twofold, significantly more than in KOR^SERT^ cKO group (Figure 1C). These data demonstrate that global deletion of KOR in SERT-expressing neurons blocks stress-induced potentiation of cocaine CPP.

**Figure 1.**
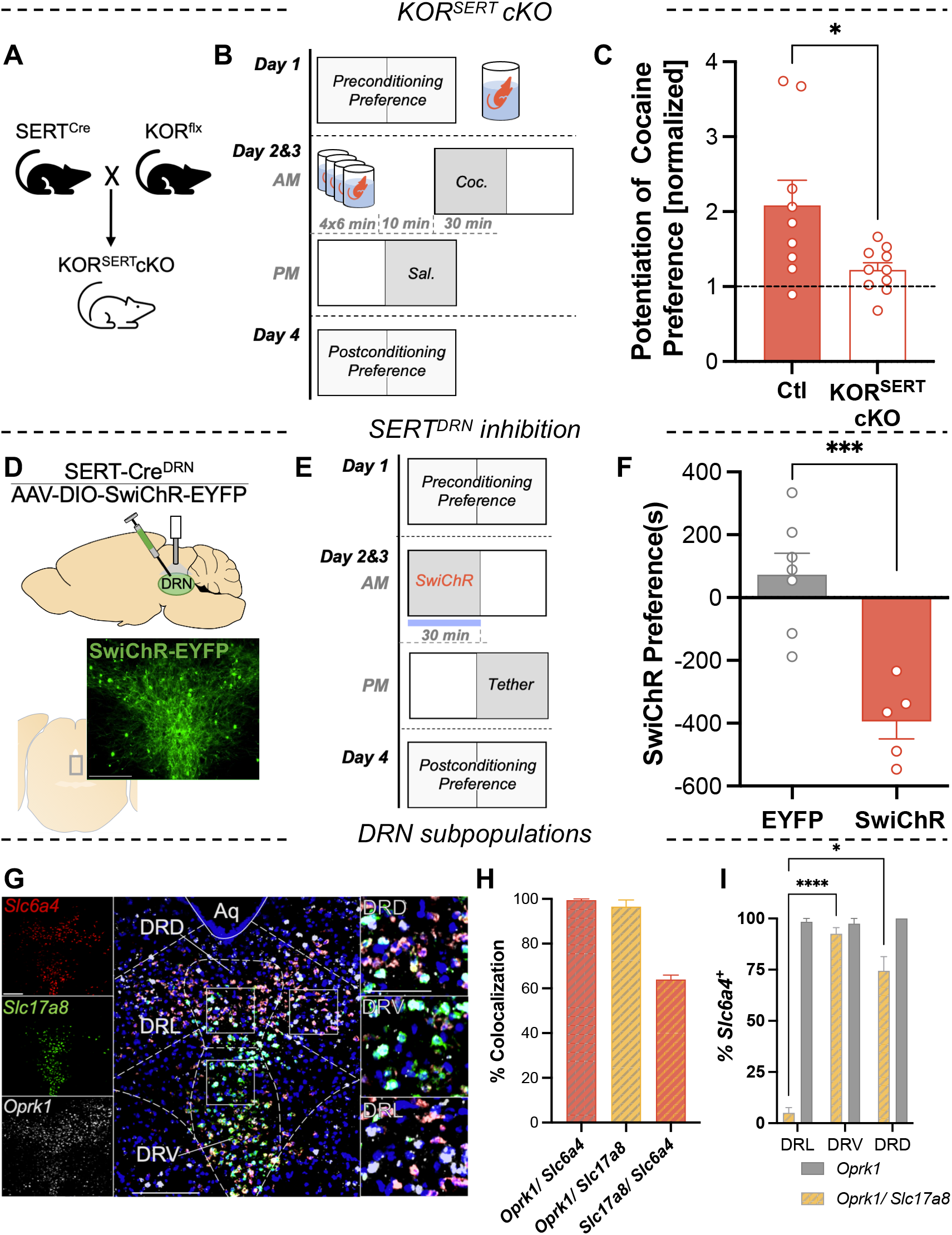
Serotonin neuron kappa opioid receptors mediate stress potentiation of cocaine reward and can be mimicked by optogenetic inhibition. (A) Breeding scheme used to excise KOR gene from SERT expressing neurons. (B) Schematic of rFSS potentiation of cocaine CPP protocol. Mice were subjected to rFSS on day 1 and day 2 prior to cocaine conditioning. (C) Cocaine preference scores with each genotype normalized to its unstressed control (n=9-10) (P= 0.036). (D) Cartoon depicting DRN injection of inhibitory opsin (AAV5-DIO-SwiChR-EYFP) and cannula placement in a SERT-cre mouse. Below: image showing EYFP expression in the DRN. Scale bar= 50 μm. (E) Schematic of optogenetic CPA protocol. During optogenetic conditioning, the mouse was confined to one chamber where it received opto-inhibition of SERT^DRN^ neurons. (F) Serotonin inhibition preference scores (postconditioning-preconditioning preference for opto-paired chamber) for control and SwiChR-inhibition conditioned groups (n=5-7) (P< 0.001). (G) Representative image showing expression of transcripts for SERT (*Slc6a4)*, VGluT3 (*Slc17a8*), and KOR (*Oprk1*) in the medial DRN. Right insets: higher magnification of rectangular regions showing colocalization in cells of the dorsal, ventral, and lateral aspects of the DRN (DRD, DRV, DRL, respectively). Scale bar= 200μm, 200μm, 25 μm. (H) Quantitation of cells co-expressing two transcripts, expressed as percentage cells in the denominator indicated (n=3) (I) Quantitation of *Slc6a4*^*+*^ cells co-expressing *Oprk1* or both *Oprk1* and *Slc17a8* in each subregion, expressed as percentage of *Slc6a4*^*+*^ cells per subregion (P< 0.001).

#### SERT^DRN^ inhibition is aversive

KOR activation by stress hyperpolarizes serotonergic neurons in the DRN [18], but stress exposure also broadly affects brain physiology. To assess the effect of selective inhibition of DRN neurons, we optogenetically inhibited SERT^DRN^ neurons by injecting AAV5-DIO-SwiChR-EYFP into the DRN of SERT-Cre mice (Figure 1D). SwiChR is a channelrhodopsin variant that conducts chloride and has been utilized to generate long-term, reversible inhibition, while avoiding photic damage [30,31]. We conducted a CPP assay by confining the mice to an optically-paired chamber during SERT^DRN^ SwiChR treatment for two days, following and preceding preference tests (Figure 1E). SERT^DRN^ SwiChR treatment induced a robust and significant aversion to the optically-paired chamber (Figure 1F). These results support the conclusion that inhibition of SERT^DRN^ neurons by either KOR activation or optogenetic inhibition produces place aversion.

#### KOR is expressed in SERT and VGluT3 DRN subpopulations

DRN serotonin neurons are anatomically and phenotypically heterogenous and have been suggested to form functional subsystems regulating diverse stress-sensitive processes [32–36], but the distribution of KOR within DRN serotonin neurons has not been evaluated. RNAscope was used to probe for co-expression of transcripts for KOR (*Oprk1)*, SERT (*Slc6a4*), and the vesicular glutamate transporter type III (VGluT3; *Slc17a8*) (Figure 1G, S1A). SERT^DRN^ and VGluT3^DRN^ neurons are both largely serotonergic populations that overlap extensively yet may have distinct roles in driving reward-related behaviors [19,20,22]. Most *Slc6a4* neurons expressed *Slc17a8*, indicating substantial overlap of SERT^DRN^ and VGluT3^DRN^ populations (Figure 1H). *Oprk1* was present in nearly all *Slc17a8*^*+*^ or *Slc6a4*^*+*^ cells (Figure 1H). The expression of KOR transcript in the majority of SERT^DRN^ and VGluT3^DRN^ cells indicates a potential for direct regulation of these subsystems by KOR.

Next, the percentage of *Slc6a4*^+^ neurons in each subregion that co-expressed *Oprk1* (*Oprk1*^+^/*Slc6a4*^+^) was determined. The percentage of *Slc6a4*^+^ neurons that co-expressed both *Oprk1* and *Slc7a18 (Oprk1*^+^/*Slc17a8*^+^/*Slc6a4*^+^) was significantly higher in the dorsal and ventral DRN than in the lateral DRN (DRL), where *Slc17a8* was nearly absent (Figure 1I). These data indicate that unlike the majority of SERT^DRN^ neurons, SERT^+^ neurons in the lateral DRN are almost exclusively *Oprk1*^+^/*Slc17a8*^−^.

#### SERT^DRN^ neurons innervate the medial NAc, but VGluT3^DRN^ neurons do not

To assess the projections of these SERT^DRN^ and VGluT3^DRN^ populations to the NAc, AAV5-DIO-ChR2-EYFP was injected into the DRN of SERT-Cre and VGluT3-Cre mice. Labeled terminals in the medial NAc (mNAc) revealed that SERT^DRN^ projection terminals in the mNAc were denser than VGluT3^DRN^ terminals, which were nearly absent (Figure 2A, 2B). These data are consistent with previous reports of different projection biases of these populations [19,32].

**Figure 2.**
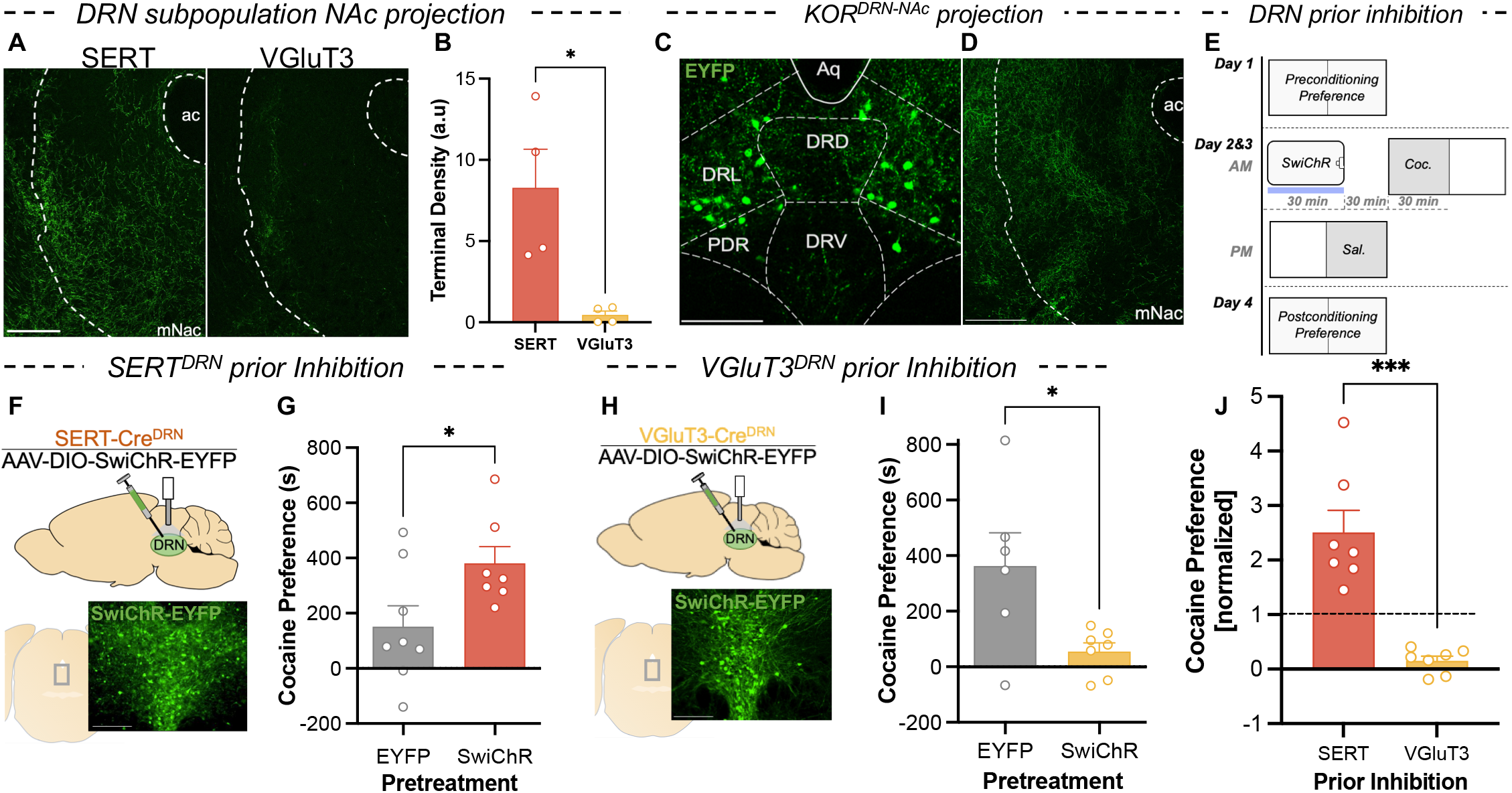
Prior inhibition of DRN serotonin subpopulations with distinct projection bias has divergent effects on cocaine preference. (A) NAc terminal expression of EYFP-tagged ChR2 in a SERT-Cre and VGluT3-Cre mouse. Scale bar=200μm. (B) Quantification of terminal density (arbitrary units) in the NAc of SERT-cre and VGluT3-cre mice (n=4). (P=0.044) (C) Expression of retrogradely-delivered EYFP in subregions of central DRN of KOR-Cre mouse. Scale bar=200μm. (D) Expression of EYFP^+^ terminals in the NAc of KOR-Cre mice. Scale bar=200μm (E) Schematic showing assay of optogenetic replication of KOR-mediated cocaine CPP potentiation. Mice received optogenetic inhibition of specified DRN subpopulations prior to cocaine conditioning. (F) Cartoon depicting injection of AAV5-DIO-SwiChR-eYFP into the DRN and placement of cannula above injection site. Below: expression of EYFP-tagged SwiChR in the DRN of a SERT-Cre mouse. Scale bar=50μm (G) Cocaine preference scores (postconditioning preference-preconditioning preference) for groups subject to control treatment and SERT inhibition prior to conditioning (n=7-8) (P= 0.037). (H) Cartoon depicting injection of AAV5-DIO-SwiChR-EYFP into the DRN and placement of the cannula above the injection site. Below: expression of EYFP-tagged SwiChR in the DRN of a VGluT3-Cre mouse. Scale bar=50μm. (G) Cocaine preference scores for groups subjected to control treatment and VGluT3 inhibition prior to conditioning (n=6-7) (P= 0.050). (H) Comparison of cocaine preference scores (normalized to respective EYFP controls) following inhibition of SERT^DRN^ or VGluT3^DRN^ neurons (n=7) (P <0.001).

#### DRN-projecting KOR neurons are restricted to the lateral DRN

To confirm that KOR is expressed within the DRN-NAc projection, a retrograde virus (AAVretro-DIO-EYFP) was injected into the NAc of KOR-Cre mice [37]. We observed a population of KOR-expressing, NAc-projecting neurons that was concentrated in the lateral aspect of DRN (Figure 2C). This indicates that DRN KOR neurons projecting to the NAc define an anatomically segregated subpopulation. To validate these findings, AAV5-DIO-ChR2-EYFP was injected into the DRN of KOR-Cre mice, and examination of the mNAc showed robust terminal expression of the fluorophore (Figure 2D). Together, these findings demonstrate that KOR is expressed in a subpopulation of DRN neurons that project to the mNAc, indicating a potential for direct regulation of this DRN-NAc projection by Dyn/KOR.

#### SERT^DRN^ inhibition recapitulates KOR-mediated potentiation of cocaine preference, but VGluT3^DRN^ inhibition does not

Although prior work has demonstrated an association between KOR activation, somatic hyperpolarization of DRN neurons, increased serotonin reuptake, and potentiation of cocaine preference, a causal link between decreased 5-HT and increased cocaine preference has not been established. We mimicked previous studies of KOR-agonist induced potentiation of cocaine preference but substituted KOR-agonist administration with optogenetic inhibition of DRN subpopulations (Figure 2E). SERT-Cre mice received DRN viral injections of AAV5-DIO-SwiChR-EYFP or AAV5-DIO-EYFP, and an optical fiber was placed above the site of viral expression (Figure 2F). These mice received 30min of SERT^DRN^ inhibition that terminated 30min prior to cocaine conditioning. Comparing cocaine preference scores following SERT^DRN^ SwiChR pretreatment to controls revealed a significant potentiation of subsequent cocaine CPP (Figure 2G). These findings indicate that prior inhibition of SERT^+^ neurons in the DRN is sufficient to potentiate cocaine CPP thereafter.

In parallel studies, VGluT3-Cre mice were injected with AAV5-DIO-SwiChR-EYFP or AAV5-DIO-EYFP in the DRN, and an optical fiber was placed above the site of viral expression (Figure 2H). Surprisingly, VGluT3^DRN^ SwiChR pretreatment significantly attenuated subsequent cocaine preference (Figure 2I). Normalizing preference scores of the SERT^DRN^ and VGluT3^DRN^ SwiChR groups to their corresponding controls illustrates divergent consequences on subsequent cocaine preference (Figure 2J). Thus, inhibition of different DRN populations exerted bidirectional control of subsequent cocaine preference, and inhibition of SERT^DRN^ neurons (but not VGluT3^DRN^ neurons) was sufficient to replicate the consequences of stress on subsequent cocaine CPP.

#### Increased 5-HT tone in the mNAc blocks KOR-mediated potentiation of cocaine preference and rFSS immobility

To directly probe the hypothesis that a KOR-mediated decrease in serotonin tone within the mNAc is necessary for potentiation of subsequent cocaine preference, we manipulated SERT^DRN-NAc^ terminals during KOR activation. SERT-Cre mice were injected AAV5-DIO-ChR2-EYFP in the DRN, and a bilateral optical fiber was placed above the mNAc (Figure 3A). One hour before each cocaine conditioning session, these mice and their controls received pretreatment that included the selective KOR agonist U50488 (5mg/kg) as well as optical stimulation of SERT terminals (to counteract KOR-induced decreases in serotonin tone) that terminated 5min before cocaine conditioning (Figure 3B). Cocaine preference scores were compared to unstressed baseline cocaine preference scores (from Figure 2G). U50488 pretreatment resulted in a potentiated cocaine preference in control animals that was attenuated by concurrent stimulation in the SERT^DRN-NAc^ ChR2 group (Figure 3C).

**Figure 3.**
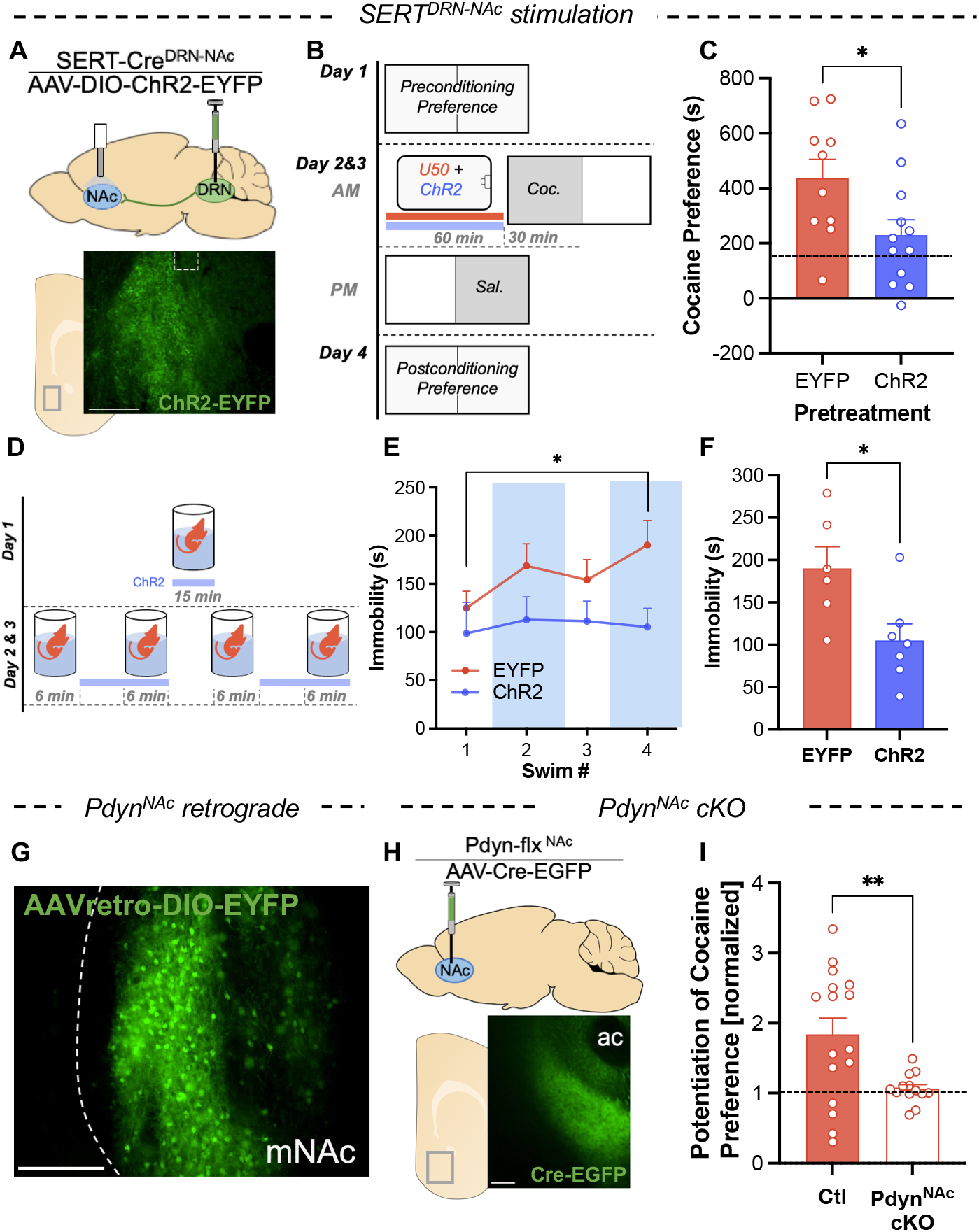
Stimulation of DRN-NAc serotonin terminals and Pdyn^NAc^ cKO blocks potentiation of cocaine preference. (A) Cartoon showing DRN injection of AAV5-DIO-ChR2-EYFP and placement of optical cannula above the NAc in a SERT-Cre mouse. Below: expression of EYFP^+^ terminals in NAc of SERT-Cre mouse. (B) Schematic of cocaine CPP with pretreatment of KOR agonist and stimulation of serotonin terminals prior to cocaine conditioning. Prior to each cocaine conditioning session, mice were pretreated with KOR agonist (U50488; 5mg/kg I.P.) and received concurrent optical stimulation of serotonin terminals in the NAc or control treatment. (C) Cocaine preference scores of mice receiving KOR agonist prior to cocaine conditioning with or without concurrent ChR2 stimulation of SERT^+^ terminals in the NAc (Dashed line shows typical unstressed cocaine preference) (n=10-12) (P = 0.029). (D) Schematic of optical stimulation of SERT terminals in the NAc during rFSS. (E) Time immobile during swim bouts on day 2 of rFSS for mice with stimulation of NAc SERT terminals and controls (n=6-7) (*Post-hoc*, P=0.038) (F) Time immobile during last swim bout of rFSS in panel E (P = 0.021). (G) Representative images showing expression of retrogradely-delivered fluorophore (EYFP) in the NAc of a Pdyn-Cre mouse. Scale bar=200μm. (H) Cartoon showing injection of virus delivering Cre-recombinase (AAV-Cre-EGFP) to the NAc of Pdyn-flx mice. Below: representative image showing expression of AAV-Cre-EGFP in the NAc. (E) Cocaine preference scores with each group normalized to its unstressed controls (n=13-16) (P=0.005).

Next, SERT^+^ terminals were stimulated during a rFSS assay to determine if decreased NAc 5-HT is required for rFSS-induced immobility (Figure 3D). Immobility was analyzed, showing an escalation of immobility in controls but not in the SERT^DRN-NAc^ ChR2 group (Figure 3E, S2A, S2B). Comparing immobility during the final swim shows that ChR2-stimulated mice spent significantly less time immobile (Figure 3F). These results suggest that stress-induced changes in mNAc serotonergic terminals are required for passive coping and potentiation of subsequent cocaine preference.

#### Pdyn^NAc^ is required for stress potentiation of cocaine preference

The role of dynorphin acting on KOR expressed in DRN-NAc projections has been demonstrated by the effects of global prodynorphin gene deletion and local KOR antagonism [1,8], but the neuronal source of dynorphin responsible for stress-induced potentiation of cocaine CPP is unknown. To identify candidate sources of dynorphin to the NAc, two different retrograde viral constructs (AAVretro-DIO-EYFP or CAV2-DIO-Zsgreen) were injected into the NAc of Pdyn-Cre mice. Examining regions for labeled neurons revealed signal only in the mNAc (Figure 3G, S2C). Distinct tropisms of these viruses have been documented that may cause either construct to undercount input populations [37]. However, because both show signal exclusively in the mNAc, we conclude that local neurons within the mNAc likely represent the principal source of endogenous dynorphin for this region.

To assess the necessity of these Pdyn^NAc^ neurons in stress potentiation of cocaine CPP, AAV5-Cre-EGFP or AAV5-EGFP was injected into the NAc of Pdyn-lox/lox (‘Pdyn-flx’) mice to generate ‘Pdyn^NAc^ cKO’ or ‘Ctl’ mice (Figure 3H). Examining the NAc of these mice demonstrated viral expression that was confined to the NAc (Figure 3I). These Pdyn^NAc^ cKO mice were subjected to a rFSS potentiation of cocaine CPP assay. Preference scores showed elevated basal preference but no rFSS potentiation of preference in the Pdyn^NAc^ cKO group (Figure S2D). To isolate the effect of stress on potentiation of cocaine preference, we normalized rFSS cocaine preference scores in each group to the cocaine preference of their unstressed counterparts (Figure 4I). This revealed a stress potentiation of cocaine preference of nearly two-fold in controls that was absent in Pdyn^NAc^cKOs. This finding supports a central role of the Pdyn^NAc^ population in regulation of both basal cocaine preference and stress potentiation of that preference.

**Figure 4.**
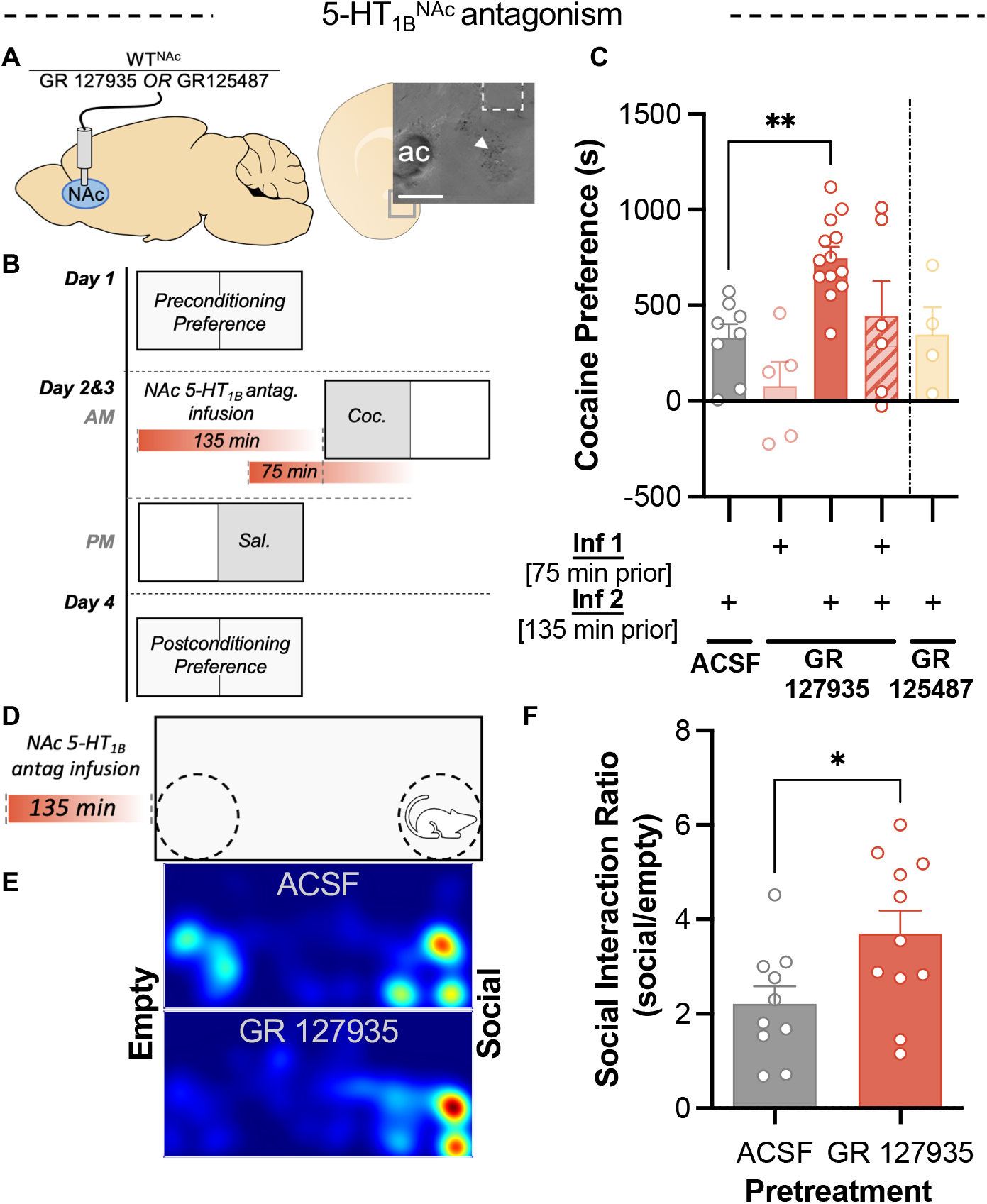
Prior 5-HT_1B_ receptor antagonism in NAc potentiates subsequent cocaine and social preference. (A) Cartoon showing implantation of fluid cannula for infusion of 5-HTR antagonists into the NAc of WT mice. (Inset) image showing dye injection (arrow) and damage from cannula confirming placement in medial NAc. Scale bar=200μm. (B) Schematic of cocaine CPP procedure showing pretreatment with NAc infusions of 5-HT_1B_ antagonist (GR 127935) at time points 75min or 135min prior to cocaine conditioning. (C) Cocaine place preference scores (postconditioning-preconditioning preference) for mice pretreated with infusions of 5-HT_1B_ antagonist GR 127935 or ACSF in the NAc (n=4-13) (*Post-hoc*, P=0.001) (D) Schematic showing pretreatment NAc infusion of 5-HT_1B_ antagonist prior to three-chamber social interaction assay. (E) Representative heatmaps indicating the distribution of time spent in the social interaction apparatus after infusion of GR 127935 into the NAc 135min prior and ACSF infusion control. (F) Social interaction ratio (time spent in social zone/time spent in empty zone) following pretreatment with GR 127935 (n=10-11) (P=0.028).

#### Blockade of mNAc 5-HT_1B_ receptors recapitulates KOR-mediated potentiation of cocaine preference and increases social preference

These finding suggest that a reduction in 5-HT tone in NAc is responsible for stress-induced potentiation of cocaine CPP, but the specific 5-HT receptor type that mediates the consequences of decreased serotonin tone is not known. Of the 5-HT receptors expressed in the NAc and known to regulate cocaine preference, the 5-HT_1B_ receptor was an especially plausible candidate [38,39]. To mimic the transient decrease in serotonin at 5-HT_1B_ receptors caused by stress, 5-HT_1B_ antagonist GR 127935 was infused into the NAc of WT mice (Figure 4A). Histological data measuring pERK-IR (Supplemental Figure 3) demonstrated that 5-HT_1B_ receptor blockade was present 75min, but not 135min, after GR 127935 infusion. GR 127935 produced a transient decrease in signaling at this receptor (Figure S3). GR 127935 was infused at 135min and/or 75min prior to each cocaine conditioning session (Figure 4B). Preference scores indicated that only infusion 135min prior to conditioning resulted in potentiation of cocaine preference (Figure 4C). Separately, infusion of a 5-HT_4_ antagonist prior to cocaine conditioning was tested and failed to potentiate preference, implying that this potentiation is not a general consequence of 5-HTR inhibition (Figure 4C). GR 127935 failed to potentiate cocaine preference in the group that received a second infusion of antagonist 75min prior to cocaine conditioning, indicating that this potentiation of cocaine CPP is sensitive to 5-HT_1B_ receptor blockade during cocaine administration (Figure 4C). These results demonstrate that a prior decrease in activation of NAc 5-HT_1B_ receptors is sufficient to potentiate subsequent cocaine preference and mimic the effects of rFSS.

To assess whether this reward potentiation is specific to cocaine or reflective of a broader change in reward processing, mice received infusions of GR 127935 or ACSF into the mNAc and 135min later were assayed for social preference, a behavior regulated NAc 5-HT_1B_ receptors [23]. In this assay, mice could explore an apparatus with an empty, inverted pencil cup in one corner and a pencil cup containing a novel mouse in the opposite corner (Figure 4D, 4E). Social interaction scores were significantly greater in mice pretreated with GR 127935 than ACSF (Figure 4F). These findings indicate that a prior blockade of 5-HT_1B_ receptors, utilized here to reflect decreased serotonin tone in the mNAc, results not only in a potentiation of cocaine reward but social reward as well.

#### rFSS increases 5-HT_1B_ transcript expression in Pdyn^NAc^ neurons

Chronic exposure to stress or psychostimulants increases 5-HT_1B_ transcript in the NAc, but regulation by sub-chronic stress exposure has not been detected [40–42]. This may be attributed to low sensitivity of prior techniques or difficulty assessing cell-type specific changes. We used RNAscope to evaluate expression of *Htr1b* in NAc subpopulations. In unstressed mice, *Htr1b* colocalized with *Pdyn, Adorsa2a,* and *Chat*-expressing cells (Figure 5A, 5B, S4). To assess the effects of stress on *Htr1b* expression, brains were dissected 30min and 24hr after rFSS and stained tissue sections were compared to unstressed controls (Figure 5C). Total levels of *Htr1b, Pdyn,* and *Adora2a* transcript did not change after rFSS (Figure 5D). In contrast, examining the levels of *Htr1b* in *Pdyn* and *Adora2a* subpopulations revealed a significant and selective increase in *Htr1b* expression in *Pdyn*^+^ cells 30min after stress (Figure 5E). These results indicate that the sub-chronic stressor (rFSS) which potentiates cocaine CPP induces a transient and selective increase in 5-HT_1B_ transcript in *Pdyn*^+^ cells of the mNAc.

**Figure 5.**
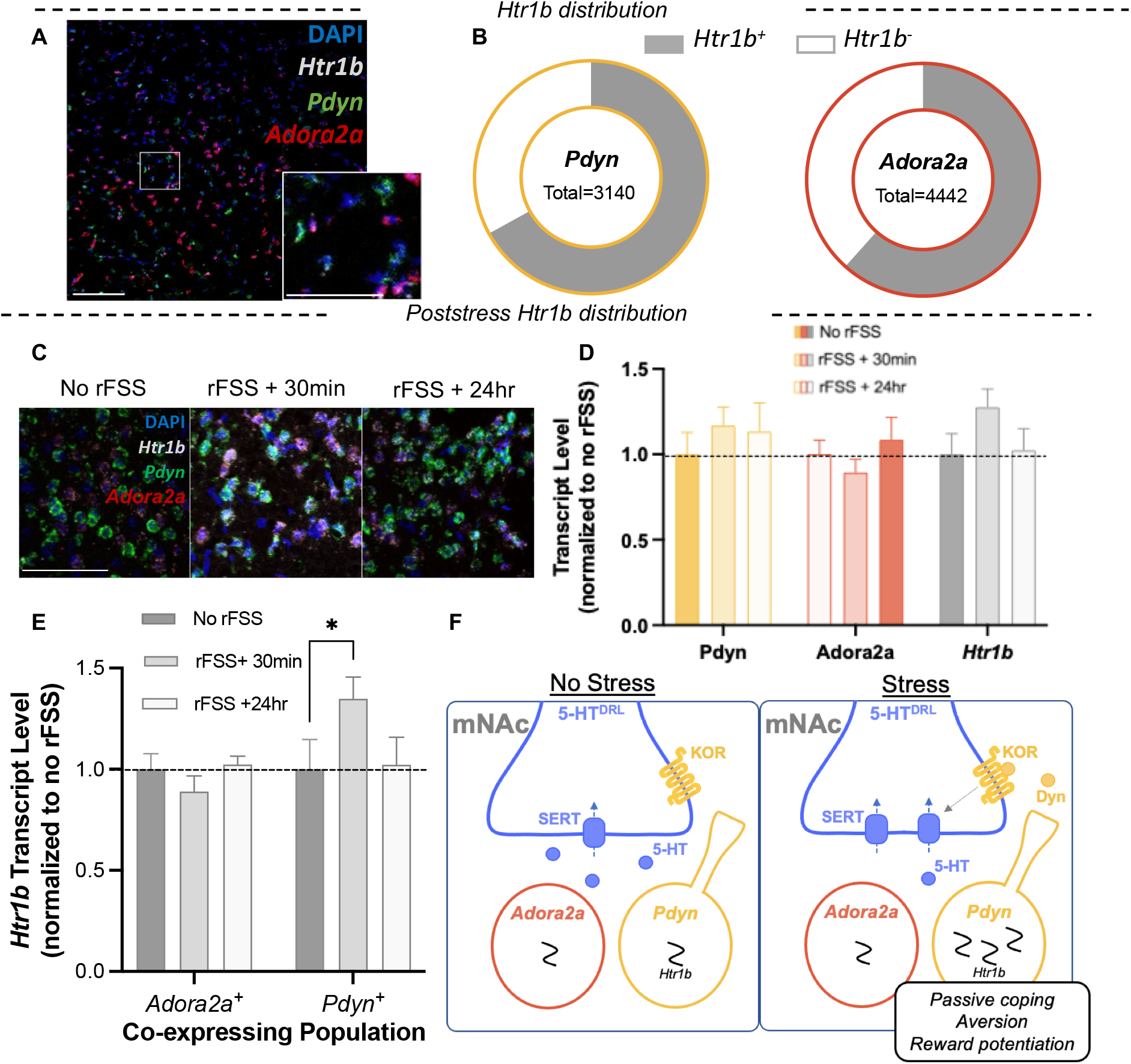
Stress induces a transient and cell-type specific increase in *Htr1b* expression in the NAc. (A) Representative image showing *in-situ* hybridization labeling *Pdyn, Adora2a, and Htr1b* transcripts in the medial NAc. Inset: higher magnification of rectangular region. Scale bar= 100μm, 25μm (inset). (B) Proportion of each subpopulation that expresses *Htr1b. (*C) Representative images showing colocalization and intensity of labelling of *Pdyn, Adora2a, and Htr1b* transcripts in the medial NAc after stress. Scale bar=25μm. (D) Quantified total levels of *Pdyn, Adora2a, and Htr1b* following stress, normalized to no rFSS controls (n=6-8). (E) Quantified levels of *Htr1b* transcripts expressed in *Pdyn*^+^ and *Adora2a*^+^ cells following stress, normalized to no rFSS controls. (*Post-hoc,* P= 0.042) (F) Summary schematic of NAc circuit mediating response to stressors, aversion, and reward potentiation. With exception of SERT, the schematic is simplified to include only entities measured or manipulated in this study.

### Discussion

The principal findings of the present study provide insight into the mechanisms by which stress impinges on the serotonin system to sensitize animals to subsequent reward. KOR expression within serotonergic neurons and dynorphin expression within the mNAc were critical to stress potentiation of cocaine reward (Figure 5F). Temporally precise manipulations of DRN circuitry showed that acute inhibition of DRN serotonin neurons drives negative affect and can potentiate subsequent cocaine preference. Additional experiments indicated that KOR-induced decreases in serotonin within the NAc control both behavioral coping and increased cocaine preference. We isolated the effect of decreased serotonin tone at the 5-HT_1B_ receptor by transient blockade of 5-HT_1B_ receptors, which was sufficient to recapitulate the potentiation of cocaine preference observed following KOR activation (Figure 5F). Lastly, we show that stress selectively increases expression of 5-HT_1B_ transcript in *Pdyn*^+^ neurons (Figure 5F). Together, this evidence details a potential Dyn-KOR-5-HT-5HT_1B_ axis contained within the mNAc, in which stress provokes a transient decrease of serotonin tone that is central to passive coping, aversion, and increased cocaine preference.

Stress and KOR activation potentiate cocaine preference through interactions at serotonin terminals in the ventral striatum, but the circuitry involved has not been fully characterized [1,5,8,17]. We indicate that the KOR-expressing neurons in the lateral DRN that project to the mNAc and Pdyn-expressing neurons located within the mNAc regulate stress potentiation of cocaine preference. Retrograde tracing of dynorphin inputs to the mNAc revealed only inputs from the mNAc itself, implying that Pdyn^NAc^ neurons may provide the sole source of dynorphin to this region. This indicates that in addition to a critical role in stress-potentiation of cocaine preference, Pdyn^NAc^ may also regulate other behaviors that are dependent on KOR activation in the NAc, including escalation of drug taking, learned helplessness, and pain-induced negative affect and anhedonia [6,20,43–46].

We observed robust aversion to SERT^DRN^ inhibition in the present study. Although place preference is not a direct measure of affect, this finding supports theories of serotonin function that assert a central role of decreased serotonin tone in negative affect [10,26]. Together with data indicating the necessity of KOR^SERT^ in stress potentiation of cocaine CPP, these results suggest that a decrease in serotonin tone may mediate stress-induced dysphoria, thereby increasing subsequent preference for cocaine. During stress potentiation of cocaine preference, stress likely decreases NAc serotonin prior to cocaine administration, but this effect is unlikely persistent, as cocaine inhibition of SERT function dramatically increases NAc serotonin tone [9,47]. Thus, temporally precise manipulation of DRN serotonin was crucial to evaluating our central hypothesis that a *prior* stress-induced decrease in serotonin tone drives subsequent increased sensitivity to drug reward. Our findings support this model and may support a general framework by which KOR and 5-HT-mediated dysphoria increases the relative impact of reward on affect.

Recently, an integral role of SERT^DRN^ neurons in regulating response to stress and reward has begun to come into focus, but how SERT^DRN^ neurons mediate the impact of stress on responses to natural and drug reward remains opaque [25,32,34,48]. We isolated the potential role of decreased serotonin tone in stress potentiation of cocaine preference first by selective inhibition of SERT^DRN^ neurons prior to cocaine conditioning and demonstrated this that a prior reduction in serotonin was sufficient to potentiate subsequent cocaine preference. Surprisingly, we found that inhibition of VGluT3^DRN^ neurons, which overlap extensively with SERT^DRN^, strongly attenuated subsequent cocaine preference. These findings further demonstrate that the DRN is a critical regulator of reward behavior, and we reason that these divergent effects may be mediated by the non-overlapping fraction of these DRN populations (i.e., SERT^+^/VGluT3^−^ neurons drive reward potentiation) and their projections. The population of overlapping VGluT3^+^/SERT^+^ DRN neurons project to a myriad of target regions, including the lateral hypothalamus, ventral tegmental area and many aspects of the cortex [33]. Based on our DRN *in-situ* hybridization and projection tracing studies, we suggest the population responsible for stress potentiation of cocaine preference may comprise *Slc6a4*^+^/*Slc17a8*^−^/*Oprk1*^+^ neurons of lateral DRN that project to the mNAc. This subregion is anatomically segregated, highly responsive to stress, and separate from parts of the DRN known to innervate other regions involved in reward processing [35,49].

Historically, the effect of stress on serotonin in the NAc has been controversial, with evidence for increases, decreases, and no effect on serotonin tone [9,50–53]. While we did not directly measure 5-HT tone, this study leveraged the neurochemically and anatomically precise nature of terminal optogenetic stimulation during stress to manipulate the SERT^DRN-NAc^ projection and indicate that a decrease in 5-HT tone within the mNAc promotes stress-induced immobility. These findings contribute to recent findings of parallel DRN serotonin subsystems innervating distinct targets to regulate reward and response to stressors [32,34]. Thus, while serotonin tone within the mNAc is a critical regulator of passive coping behavior, serotonin tone in other regions may also regulate passive coping, possibly in coordination with the projection we detail.

In this study, we examined potential contributions of the 5-HT_1B_ receptor to stress potentiation of reward. 5-HT_1B_ receptors are Gi-coupled GPCRs that inhibit neurotransmitter release and have been implicated in regulating the response to stressors and psychostimulants [39]. However, the direction of these effects is dependent on brain region, involvement of autoreceptors or heteroreceptors, and stage of addiction cycle [54–57]. In the mNAc, transient overexpression of 5-HT_1B_ heteroreceptors during mild stress enhances stress-induced potentiation of the psychomotor effects of amphetamine [39]. We observed that transient antagonism of 5-HT_1B_ receptors in the NAc was sufficient to recapitulate potentiation of cocaine preference induced by stress or decreased serotonin tone. Transient antagonism of NAc 5-HT_1B_ receptors also potentiated social preference, a behavior mediated by NAc 5-HT_1B_ receptors [23]. Together, these data suggest that 5-HT_1B_ receptors in the NAc act as a critical signal transducer, sensing a decrease in 5-HT tone and initiating postsynaptic consequences that result in potentiation of reward. Our findings also indicate that the mechanism by which decreased 5-HT_1B_ activation results in potentiation of subsequent cocaine reward may generalize to processing of other rewarding stimuli, including natural rewards. Lastly, we showed that potentiation of cocaine preference induced by transient prior blockade of NAc 5-HT_1B_ was attenuated by an additional antagonist infusion that blocked 5-HT_1B_ receptors during cocaine conditioning. These findings indicate that the 5-HT_1B_ receptor is not only involved in initiating reward potentiation but is involved in mediating the expression of increased reward as well.

5-HT_1B_ transcript is highly expressed in the NAc, and prior work indicates that chronic exposure to stressors or psychostimulants may regulate its expression within the accumbens [40,41]. Colocalization of the 5-HT_1B_ transcript *(Htr1b)* with markers of the direct pathway (*Pdyn*), indirect pathway (*Adora2a),* and cholinergic interneurons (*Chat)* showed uniform distribution across these cell types, indicating that serotonin actions through 5-HT_1B_ receptors may regulate these populations in concert to modulate processing in the NAc. Following stress, 5-HT_1B_ mRNA increased within *Pdyn*^+^, but not *Adora2a*^+^ neurons, suggesting that stress selectively increases the expression of 5-HT_1B_ transcript in cells of the direct pathway. Whether this increase in transcript is accompanied by an increase in functional receptors remains to be tested. Prior work has also shown that overexpression of 5-HT_1B_ receptors in the NAc sensitizes rats to the hedonic properties of cocaine [38,58], which intimates that stress-induced increases of postsynaptic 5-HT_1B_ receptors in direct pathway neurons may mediate stress potentiation of reward.

Chronic stress induces hedonic and motivational deficits that contribute to depression-like behavior, but sub-chronic stress can provoke coping responses. This coping response may include a proadaptive hedonic allostasis that increases sensitivity to reward and is reflected by increases in mNAc 5-HT_1B_ expression. We indicate this coping response is maladaptive in the context of drug exposure, resulting in increased drug preference and enhanced addiction risk. While our data are consistent with this interpretation, future work is required to assess the behavioral and cellular tenets of this theory. In human studies, polymorphisms of the 5-HT_1B_ receptor have been associated with major depression and substance use disorder [59–61], and receptor binding studies show altered 5-HT_1B_ binding in the NAc of subjects with major depression and alcohol dependence [62,63]. Such findings suggest a central role of NAc 5-HT_1B_ receptors in regulation of affect and substance use, but a potential connection to the Dyn/KOR system and stress potentiation of addiction risk has not been previously evaluated. The insights gleaned from this study support a functional Dyn-KOR-5-HT-5-HT_1B_ axis in which decreased 5-HT is a central regulator of drug preference, affect, and response to stressors. Future studies will be required to directly evaluate the consequences and kinetics of the stress and dynorphin-mediated effects on functional 5-HT_1B_ receptors and evaluate the therapeutic potential of this dynamic circuit.

## Abbreviations

KOR: Kappa opioid receptors
SERT: plasma membrane serotonin transporter
CPP: conditioned place preference
DRN: dorsal raphe nucleus
*Slc17a8*, VGluT3: vesicular glutamate transporter III
mNAc: medial nucleus accumbens
ac: anterior commissure
Pdyn: prodynorphin gene (*Pdyn*) or peptide
Dyn: dynorphins
*Adora2a*^+^: adenosine A_2A_ receptor
*ChAT*^+^: choline acetyltransferase
CRF: corticotropin-releasing factor
5-HT: serotonin
norBNI: norbinaltorphimine-HCl
ACSF: artificial cerebrospinal fluid
AAV: adeno-associated virus
cKO: conditional gene knockout
rFSS: repeated forced swim stress
WT: wild-type mice
*SERT*^*DRN-NAc*^: SERT^+^ neurons projecting from DRN to NAc
−IR: immunoreactivity
PBS: phospho-buffered saline
ISH: fluorescent in situ hybridization
SERT^DRN^: SERT^+^ neurons within the DRN
VGluT3^DRN^: VGluT3^+^ neurons within the DRN
Pdyn^NAc^: Pdyn^+^ neurons within the NAc
KOR^flx^: KOR-lox/lox
Pdyn^flx^: Pdyn-lox/lox

## Funding and Disclosure

This research was supported by USPHS grants P50-MH106428 (CC), T32GM007750 (HMF), R01-DA030074 (CC), R01 DA041356 (JN), and S10 OD016240 (Keck Center). The authors have no conflicts of interest to disclose.

## Acknowledgements

We would like to thank Drs. Larry Zweifel, Sarah Ross, and Richard Palmiter for providing reagents; Dr. Kevin Coffey for assistance with MATLAB scripts used in RNAscope analysis; Zeena Rivera for genotyping assistance; and the WM Keck Microscopy Center for microscopy support.

## Author contributions

HMF, PS, CN, RT and AA conducted the experiments. HMF, PS, RT analyzed the results. All of the authors contributed to the design of the experiments. HMF and CC wrote the manuscript. JFN, PS, and BBL provided editorial comments.

## Supplemental Materials and Methods

### Drugs

Cocaine-HCl, norbinaltorphimine-HCl (norBNI), and ±U50488 were provided by the National Institute on Drug Abuse Drug Supply Program (Bethesda, MD) and were dissolved in 0.9% saline. Sodium pentobarbital, Beuthanasia Special-D, and isoflurane were obtained from University of Washington Medical Center Drug Services. CP 94253-HCl, GR 127935-HCl, and GR 125487 were purchased from Tocris Bioscience and dissolved in artificial cerebrospinal fluid (ACSF).

### Viral reagents

CAV2-DIO-ZsGreen was provided by Dr. Larry Zweifel (University of Washington). UNC Vector Core or Addgene provided: AAV5-DIO-EYFP (UNC/ Addgene #27056), AAV5-DIO-SwiChR_CA_-EYFP (UNC), AAV5-DIO-ChR2-EYFP (UNC/ Addgene #20298), AAV5-EGFP (#105547), AAV5-Cre-EGFP (Addgene #105545), and AAVrg-DIO-EYFP (Addgene #27056). Viral suspensions were stored at −80°C until use and injected undiluted (2×10^12^ - 3×10^13^ vg/ml).

### Animals

Adult (8-20wk) male C57BL/6 mice and transgenic strains on C57BL/6 genetic background were group housed (2-5/cage), given access to food pellets and water *ad libitum*, and maintained on a 12hr light:dark cycle (lights on at 7AM). All animal procedures were approved by the University of Washington Institutional Animal Care and Use Committee and conformed to US National Institutes of Health guidelines. We obtained *Slc6a4*-Cre (SERT-Cre) mice from the GENSAT project (MMRRC:017260-UCD), *Oprk1*-Cre (KOR-Cre) mice from Dr. Sarah Ross (University of Pittsburgh) [1], *Pdyn-*IRES-Cre (Pdyn-Cre) and *Pdyn*-lox/lox (Pdyn-flx) mice from Dr. Richard Palmiter (University of Washington), *Slc17a8*-Cre (VGluT3-Cre)and *Oprk1*-lox/lox (KOR-flx) from Jackson Labs (MGI:5823257, MGI:5316477). KOR^SERT^ conditional knockout (cKO) mice were generated as previously described [2].

### General behavioral methods

Mice were kept in the same housing facility in which behaviors were assayed for at least 1 week prior to experimentation. For all optogenetic experiments, controls were Cre^+^ mice injected with AAV-DIO-EYFP instead of the active opsin. Cage changes were conducted no less than 3 days prior to behavioral testing to minimize confounding effects of environmental stress exposure. Mice were habituated to handling daily for 3 days prior to the initiation of each experiment. All experiments were conducted on mice naïve to prior treatment, except for the optical stimulation during rFSS and social approach assays, which were conducted 2 weeks after the completion of cocaine CPP. EthoVision Software (Version 3.0 & 11.0, Noldus Information Technology) was used to assess movement and generate path heatmap graphics. Experiments were conducted in sound-attenuating behavioral rooms with medium-intensity lighting.

### Stereotaxic Surgery

For aseptic surgery, mice were anesthetized in an induction chamber with 4% isoflurane before placement into a stereotaxic frame (David Kopf Instruments Model 1900) where they received 1-2% isoflurane as described previously [2]. Viral injections were performed using Hamilton Neuros syringe (Sigma-Aldrich) at a rate of 100nl/min (500nl for all behavioral studies, 750nl for tracing studies). The syringe was left in place for 5min following the injection. Injection sites were as follows: DRN (AP −4.35, ML 0, DV-2.7; 20° angle) or NAc (AP+1.35, ML +−0.7, DV −4.6) and optic fibers (Doric) were placed 0.5mm above the target site. All NAc viral injections for behavioral studies and optical stimulation were bilateral and unilateral for retrograde tracing. For drug microinfusion, guide cannula (Plastics One #C235G/SPC-1.4mm) were placed above the NAc (AP +1.35, ML +−0.7, DV −4.1), with internal cannula projecting 0.5mm past the guide cannula. Implants were secured using Metabond (Parkell) and dental cement (Stoelting). Following surgeries, mice were given rimadyl for 5 days to reduce inflammation and pain and allowed time for recovery and viral expression (10 days for infusion studies, 4 weeks for somatic optical stimulation, and 5 weeks for terminal optical stimulation and anatomical tracing studies).

### Forced Swim Stress

Mice were subjected to a modified Porsolt forced swim stress (rFSS) as described previously [3]. All swim sessions were conducted in 31⩲1°C water. On day 1, mice received a 15min initial swim, followed 22hr later by four 6min swims, each separated by 6min. After each swim, mice were removed from the water, towel dried, and returned to their home cage.

### Optogenetic stimulation during rFSS

Mice were connected to the optical tether 1min prior to swim sessions on day 1 and 2, and they remained tethered throughout the swim session. Mice were visually monitored during swim sessions. Mice that submerged due to impaired swimming were removed from the water and excluded from subsequent analysis (3 EYFP-injected and 2 ChR2-injected mice were excluded). Optical stimulation was delivered 1min prior to the initial swim and 6min prior to the second and fourth swim on the day 2.

### Cocaine conditioned place preference

Mice were assayed in a balanced place conditioning apparatus with distinct visual and tactile cues in each chamber as previously described [2,4]. On day 1, an initial preference test was conducted for place preference bias. Conditioning occurred on days 2 and 3, consisting of cocaine administration and 30min confinement to the drug-paired chamber in the morning (15mg/kg, IP) and saline administration (10ml/kg, IP) and confinement to the other chamber 4hr later. On day 4, mice were allowed to freely explore the apparatus for a postconditioning assessment in the absence of drug. Preference tests and conditioning sessions lasted for 30min and were conducted in sound attenuating chambers.

### Manipulations prior to cocaine conditioning

#### Repeated forced swim stress prior to cocaine conditioning

mice were subjected to rFSS (as described above) 30min after the initial preference test on day 1 and before cocaine conditioning on day 2, terminating 10min prior to cocaine administration.

#### Optogenetic inhibition of DRN subtypes prior to cocaine conditioning

VGluT3-Cre and SERT-Cre mice expressing EYFP or SwiChR were tethered to fiber optic cables coupled to a 473-nm laser in an empty cage bottom and received optical stimulation (0.33 Hz, 15ms pulse duration) for 30min on each conditioning day. Following optical stimulation, mice were returned to their home-cage 30min prior to each cocaine conditioning session.

#### Optogenetic excitation of SERT^DRN-NAc^ terminals during U50488 pretreatment

mice received the selective KOR agonist U50488 (5mg/kg, IP) 1hr prior to cocaine conditioning and were immediately tethered to optical fibers providing 473-nm stimulation (15Hz, 10ms pulse duration) in an empty cage bottom. Mice were untethered 5min before each cocaine conditioning session.

#### Local infusion of 5-HT receptor antagonists prior to cocaine conditioning

Wild-type (WT) mice with guide cannula placed in the NAc received infusions of the 5-HT_1B_ antagonist GR 127935 or 5-HT_4_ antagonist GR 125487 (1μg/0.2μl in ACSF, 0.1μl/min) 135min and/or 75min prior to each cocaine conditioning session. Following infusions, mice were returned to their home cage.

### Conditioned place aversion

Cannulated SERT-Cre mice expressing EYFP or SwiChR were assayed in a balanced place conditioning apparatus with distinct visual and tactile cues as previously described [2]. An initial preference test was performed on day 1 to assess baseline preference. On days 2 & 3 optogenetic conditioning was performed, comprising a tethering session with confinement to the less-preferred chamber in the morning and optical stimulation (0.33 Hz, 15ms pulse duration) with confinement to the more-preferred chamber 4hr later. On the 4^th^ day, mice were allowed to freely explore the apparatus for a final preference test in the absence optical stimulation.

### Social approach

Social interaction was assessed as described previously using a three chambered apparatus with two clear internal partitions [5]. The day prior to the experiment, age-matched target mice were habituated to confinement in an inverted pencil cup (Spectrum Diversified Designs) for 1hr. WT mice with cannula placed in the NAc received infusions of 5-HT_1B_ antagonist GR 127935 (1μg/0.2μl in ACSF; 0.1 μl/min), and 125min later were allowed to freely explore the social interaction apparatus for a 10min habituation period. The mouse was then briefly removed to a holding cage, and two inverted pencil cups were placed in the far corners of the apparatus, with one cup containing a target mouse. The experimental mouse was then reintroduced and allowed to explore for an additional 10min. Time spent in an interaction zone adjacent to each cup was recorded.

### Local infusion of 5-HT_1B_ ligands prior to histology

WT mice with guide cannula placed in the NAc received norBNI (10mg/kg, IP) 24hr prior to the experiment to minimize the effects of infusion-induced stress on pERK-IR [6]. Mice received control infusions (ACSF, 0.2μl) in the right hemisphere and drug infusions (GR 127935 or 5 CP94253; 1μg/0.2μl, 0.1μl/min) in the left hemisphere. Drug infusions consisted of CP 94253 alone or following infusions of GR127935 135min or 75min prior. 15min after infusion of CP 94253, mice were deeply anesthetized, transcardially perfused, and brains were prepared for histology as described below.

### Immunohistochemistry

Mice were transcardially perfused with 4% paraformaldehyde in 0.1M phospho-buffered saline (PBS) as reported previously [7]. Brains were then dissected, cryoprotected with 30% sucrose at 4°C overnight, frozen, cut into 40μm sections (Leica microtome, SM200R), and stored in 0.1M PBS w 0.1% sodium azide at 4°C until further processing. Standard immunohistochemical procedures were used to stain NAc sections as described previously [6]. Briefly: floating sections were washed 3×5min in PBS, then blocked for 1hr in 5% normal goat serum (Vector Labs), 0.3% Triton-X in PBS before 24hr at room temperature incubation with primary antibodies: 1:400 rabbit anti-pERK antibody (CS4370, Cell Signaling) for phospho-ERK detection or 1:1000 Chicken anti-GFP (AB12970, Abcam) to enhance detection of anterograde and retrograde tracing. Sections were washed again 4×5min in PBS before incubation with the 1:500 goat anti-rabbit 488 or goat anti-chicken 488 (Life Technologies) for 2hr at room temperature. Lastly, sections were washed 4×5 in PBS, then once with 0.5X PBS before mounting on Fisher Superfrost slides (Sigma-Aldrich) and coverslipped using Vectashield (Vector Laboratories).

### Fluorescent in situ hybridization (ISH) using RNAscope

Brains were rapidly dissected and flash frozen on dry ice. For stress experiments, brain dissection was performed either 30min or 24hr following the last swim session of the rFSS protocol or unhandled (no rFSS) controls. Thin (14μm) coronal sections containing the NAc or DRN were collected and mounted onto Superfrost plus slides using a cryostat (Leica CM 1850) maintained at − 20°C. RNAscope ISH was performed according to the Advanced Cell Diagnostics as previously reported [8]. Each set of staining included a negative control, in which probes were omitted from the process. Probes were discriminated using tyramide signal amplification (TSA) fluorophores (NEL744001, NEL745001, NEL741001; Akoya Biosciences).

#### Characterization of DRN subpopulations

Probes for *Oprk1 (mm-Oprk1), Slc6a4 (mm-Slc6a4), and Slc17a8 (mm-Slc17a8)* were used to label tissue from the central DRN (AP +4.3−4.5) of unstressed mice*. NAc Htr1b distribution:* sections containing the central NAc (AP 1.1-1.3) were obtained from stressed and unstressed mice, and separate sets of tissue were stained with probes to *Htr1b(mm-Htr1b)*/*Chat (mm-Chat)* and *Htr1b/Pdyn (mm-Pdyn)/Adora2a (mm-Adora2a).*

### Microscopy and Image Quantification

All images used for quantitation were taken using a confocal microscope (SP8X, Leica Microsystems), except for initial determination of expression in retrograde tracing studies and confirmation of viral expression in behavioral studies, in which a scanning widefield microscope was used (DMI6000, Leica Microsystems).

#### Anterograde tracing

Two brain sections containing the NAc (AP +1.3 and AP +1.0) were imaged at 10x magnification for each SERT^DRN^ ChR2 and VGluT3^DRN^ ChR2 mouse. Boundaries of the mNAc were determined using the Paxinos atlas [9]. Terminal density was determined using Image J software (NIH) by binarizing the images and calculating the density of positive pixels within the mNAc as described previously [10].

#### Retrograde tracing

For KOR retrograde tracing from the mNAc, every sixth section within the DRN (AP −4.0 to AP −5.0) was mounted and imaged at 10x magnification following anti-GFP staining to enhance detection of labeled cell bodies. For Pdyn retrograde tracing from the mNAc, every 12^th^ section throughout the collected tissue (AP +2.5 to AP −5.5) was mounted to survey across all brain regions and imaged at 10x on a scanning widefield microscope. These were visually inspected for signal by an observer naïve to treatment. Subsequently, confocal images were taken of the medial NAc at 10x magnification.

#### Effects of 5-HT_1B_ drug infusion on pERK in the NAc

The left and right medial NAc of two sections (AP +1.3 and AP +1.0) were imaged for each animal. Z-stacks (5μm thick, 7 steps) were taken at 60x magnification and an average projection was generated using LASX software (Leica Microsystems). A detection threshold for each section was set according to the brightest 5% of pixels in the ACSF image and positive cells (more than half of cell above threshold) for each image were quantified manually by an observer blind to treatment using ImageJ software (NIH). Average counts of two sections for the ACSF and drug treated hemispheres were taken for each animal. Percent increase of NAc pERK-IR^+^ cells in the drug-treated hemisphere was calculated as (pERK_Drug_-pERK_ACSF_)/ pERK_ACSF_.

#### Fluorescent ISH: DRN subpopulations

The entire DRN was imaged at 20x magnification from one section (AP 4.4) of each subject. Images were taken during the same imaging session, and capture settings were adjusted such that no signal was observed in negative controls and kept constant for all subsequent images. *NAc Htr1b distribution:* a region immediately medial and ventral to the anterior commissure (ac) was imaged at 20x magnification, with capture settings adjusted to ensure no signal in the negative controls. Two bilateral images from two sections were taken. Images were processed using custom MATLAB scripts for positive cells, co-expressing cells, and levels of RNA detected per cell; these values were averaged for each subject animal and then group averages were calculated. For all analyses, total cells were determined by the number of DAPI-stained nuclei.

### Data Analysis

Sample sizes were based on prior studies but were not predetermined by statistical methods. All aspects of histology and histological analysis were performed by an experimenter blind to genotype and treatment. Prior to further analysis, outliers in data sets were excluded using Grubb’s Test for statistical outliers. The assumption of normal distribution was tested for each data set and was statistically corrected for when this criterion was not met. *Post-hoc* tests used were Sidak’s or Dunnett’s where appropriate, with *α*=0.05.

## Supplemental Results

### Supplemental to Text Figure 1

Consistent with previous reports [3], unstressed control mice developed significant cocaine place preference, and rFSS induced robust potentiation of CPP (Figure S1A). Cocaine place preference of the unstressed KOR^SERT^cKO mice was not significantly different from that of littermate controls (two-way ANOVA; F_1, 34_= 0.27, P= 0.604), indicating that excision of KOR in SERT-expressing neurons does not regulate basal cocaine preference (Figure S1A). There was a significant main effect of stress (F_1, 34_= 8.87, P=0.005) and a marginal interaction (gene X stress interaction, F_1, 34_= 3.45, P= 0.072). Comparison within each genotype revealed a significant effect of stress in controls (Sidak *post-hoc*, P= 0.004) that was absent in the KOR^SERT^ cKO group (P= 0.664) (Figure S1A).

We found that KOR transcript was expressed in the majority of DRN cells (84±8.0%) (Figure S1B). *Slc6a4* and *Slc17a8* were present in a roughly equal percentage of DRN neurons (35±9.9% and 31±7.5%, respectively).

**Supplemental Figure S1.**
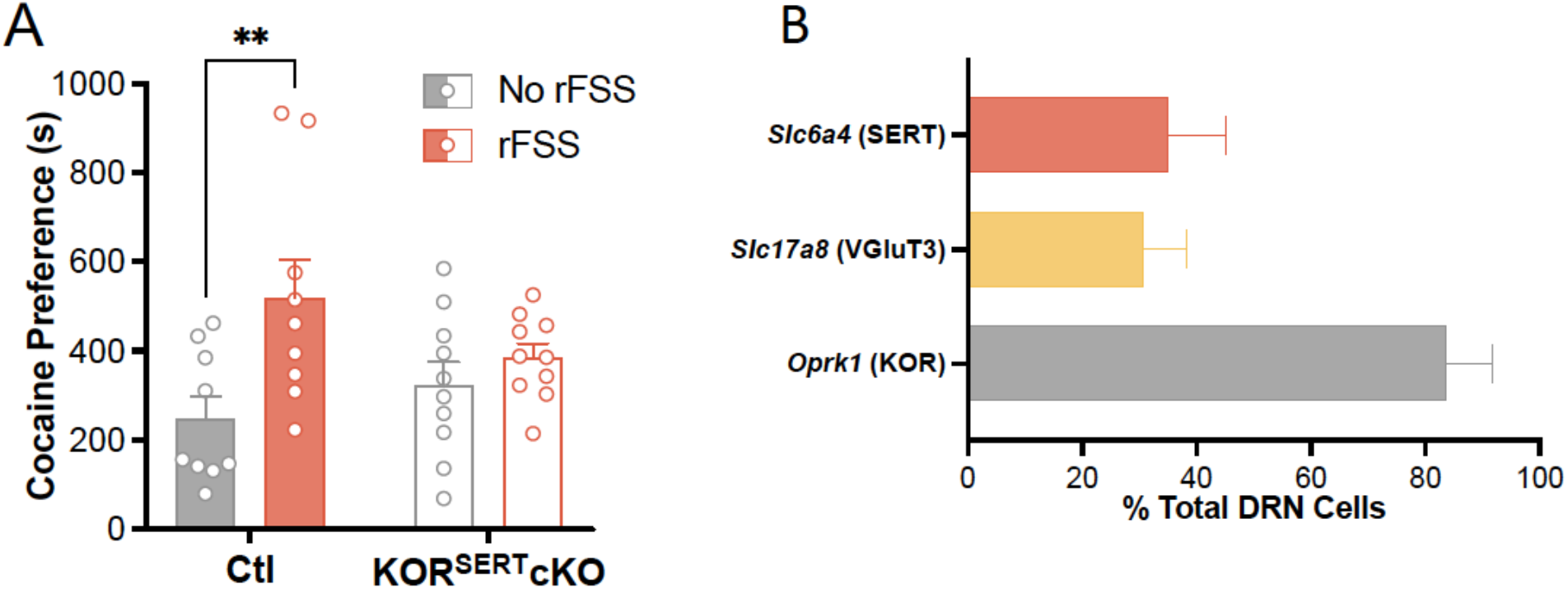
(A) Cocaine preference scores (postconditioning preference- preconditioning preference for drug-paired chamber) for control and KOR^SERT^ cKO mice with or without prior stress (n=9-10). (B) Quantitation of cells expressing transcripts for SERT, VGluT3, KOR, expressed as percentage of total DRN cells (n=3).

### Supplemental to Text Figure 3

Movement was tracked during swim sessions and indicated a decrease in movement in the EYFP group that was not apparent in SERT^DRN-NAc^ ChR2 group (Figure S2A). To calculate the escalation of immobility in mice of each group, time immobile during first swim was subtracted from the last swim of day 2 (Figure S2B). The results indicate that stimulation of serotonergic terminals in the NAc prevents escalation of immobility within subjects (unpaired, two-tailed t test, P =0.014).

Cocaine preference scores indicated a main effect of Pdyn^NAc^ excision (two-way ANOVA, F_1, 54_= 4.73, P= 0.034), indicating that dynorphin within the NAc regulates basal cocaine preference (Figure S2D). There was also a significant main effect of stress (F_1, 54_= 13.1, P= 0.001) but no significant interaction (F_1, 54_= 2.44, P= 0.12). There was a significant potentiation of cocaine preference in Cre^−^ controls (Sidak *post-hoc*, P= 0.015) but not in the Pdyn^NAc^cKOs (P=0.90). We have routinely observed cocaine preference scores higher than the ~500 s preference in the unstressed Pdyn^NAc^cKO group, indicating that the lack of stress potentiation is likely not due to a ceiling effect on expressed preference.

**Supplemental Figure S2.**
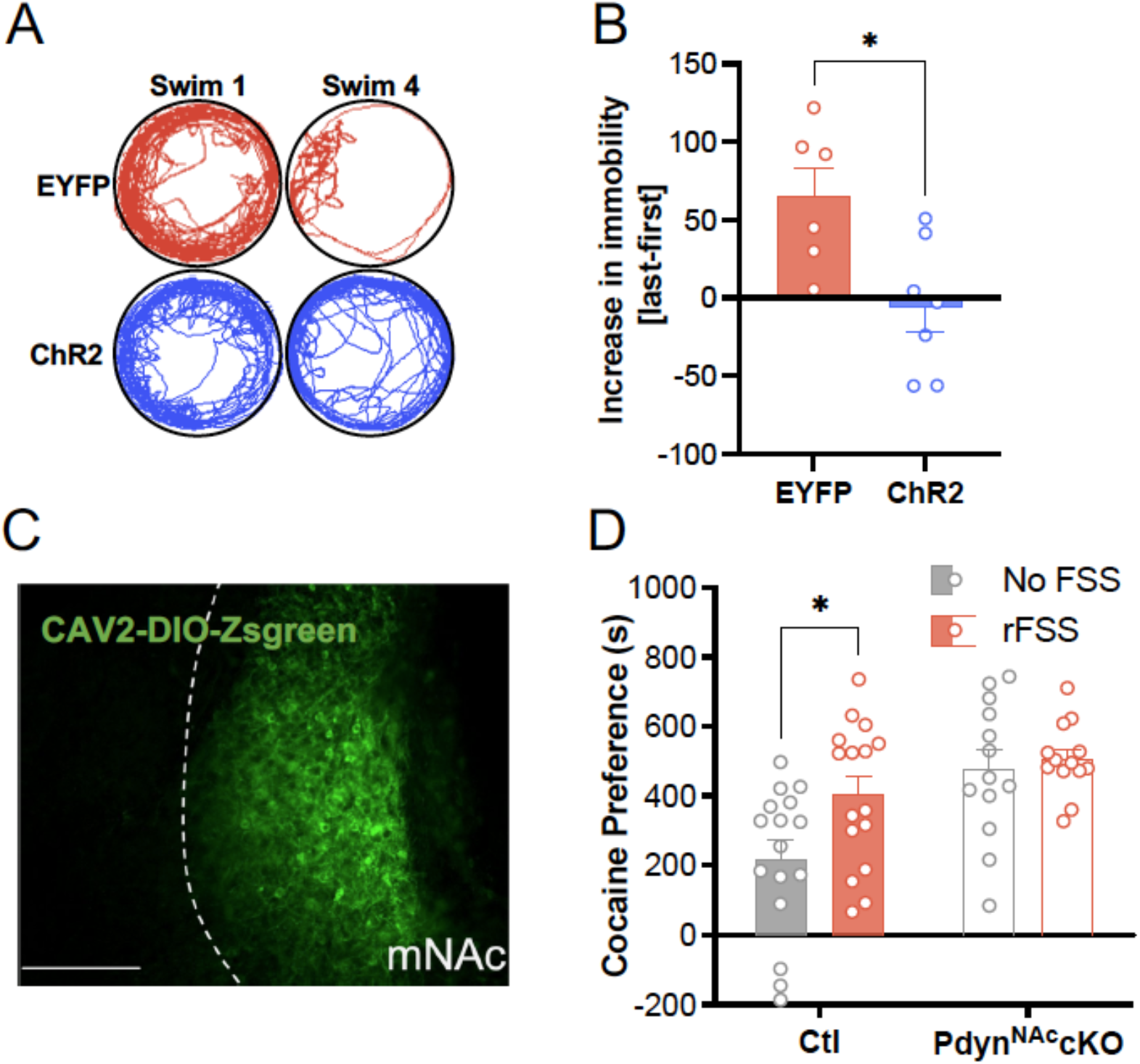
(A) Representative track traces during first and last swims on day 2. (B) Change in immobility from first to last swim on day 2 (n=6-7). (C) Representative image of ZsGreen expression in NAc of a Pdyn-Cre mouse. (D) Raw cocaine preference scores for control and Pdyn^NAc^ cKO mice with or without prior stress (n=13-16).

### Supplemental to Text Figure 4

Two time points were selected to evaluate the duration of antagonist-mediated blockade of the receptor. Injection cannula were placed in the mNAc of WT mice and 5-HT_1B_ antagonist GR 127935 was infused unilaterally (Figure S3A). This was followed either 75min or 135min later by unilateral infusion of 5-HT_1B_ agonist CP 93129 and perfusion of the brain 15min later. ACSF was infused in the opposite hemisphere during each drug infusion to control for nonspecific effects of the infusion procedure on ERK phosphorylation. Coronal sections containing the mNAc were probed for pERK-immunoreactivity (IR), a consequence of 5-HT_1B_ activation (Figure S3B) [11]. Comparison of treatments showed a significant main effect of agonist treatment on agonist-stimulated pERK-IR and a significant interaction (two-way ANOVA, agonist main effect, F_1,25_ = 7.12, P= 0.013; pretreatment X agonist interaction, F_2,25_= 8.04, P= 0.002). *Post-hoc* comparisons indicated that treatment with CP 93129 induced a significant increase in pERK-IR cells as compared to the ACSF control hemisphere in the absence of antagonist pretreatment (Sidak *post-hoc, P=* 0.002). This result confirmed that local infusion of the 5-HT_1B_ agonist induces pERK-IR within mNAc cell bodies. Pretreatment with the antagonist GR 127935 75 min prior to CP 93129 infusion blocked the agonist-induced increase in pERK-IR (Sidak *post-hoc*, P= 0.293), whereas pretreatment 135min prior did not (Sidak *post-hoc*, P= 0.033) (Figure S3C). To further interrogate antagonist duration, the number of pERK+ cells were normalized to the ACSF hemisphere for each subject, and this value was expressed as a fraction of the effect from agonist treatment alone (Figure S3D). This direct comparison of antagonist pretreatment timing illustrates the transient block of pERK-IR caused by GR 127935 infused 75min before CP 93129, but not 135min before agonist (P= 0.014).

**Supplemental Figure S3.**
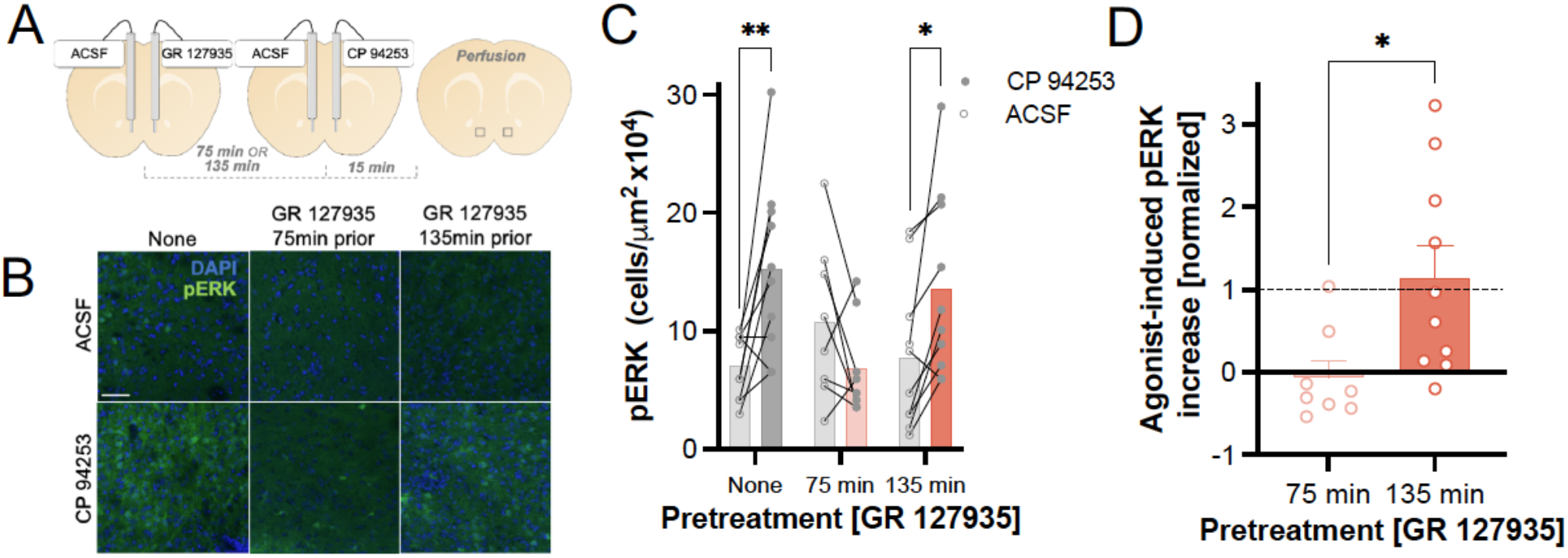
(A) Schematic of unilateral NAc infusion of 5-HT_1B_ antagonist GR 127935, followed 75 or 135min later by infusion of 5-HT_1B_ agonist CP 94253 and perfusion. Rectangles show imaging field. (B) Representative images showing immunofluorescent lab phospho-ERK1/2 (pERK) in ACSF (control) hemisphere and hemisphere receiving 5-HT_1B_ ligand infusions. Scale bar= 50μm. (C) Quantification of pERK+ cells in control hemisphere and agonist-infused hemispheres following NAc infusion of GR 127935 prior to CP 94253 (n=8-10). (D) Quantification of pERK+ cells following GR 127935 pretreatment, expressed as percentage of agonist-induced increase in pERK compared to ACSF and normalized to agonist-induced increase without pretreatment (n=8-10).

### Supplemental to Text Figure 5

Staining and imaging were performed to assess the colocalization of *Htr1b* with *Pdyn* and *Adora2a*. Results indicate that nearly half (43±2.7%) of the neurons within the mNAc express *Htr1b* (Figure S4A). Staining for other markers show that *Pdyn* and *Adora2a* were detected in a sizeable fraction of neurons (33±2.5% and 45±2.5%, respectively). A second experiment was performed to establish *Htr1b* colocalization with *Chat* (Figure S4B, S4C). *Chat* was expressed sparsely (2.5±0.5%), in line with prior findings, and *Htr1b* was present in a majority of these neurons (Figure S4C, S4D) [12].

**Supplemental Figure S4.**
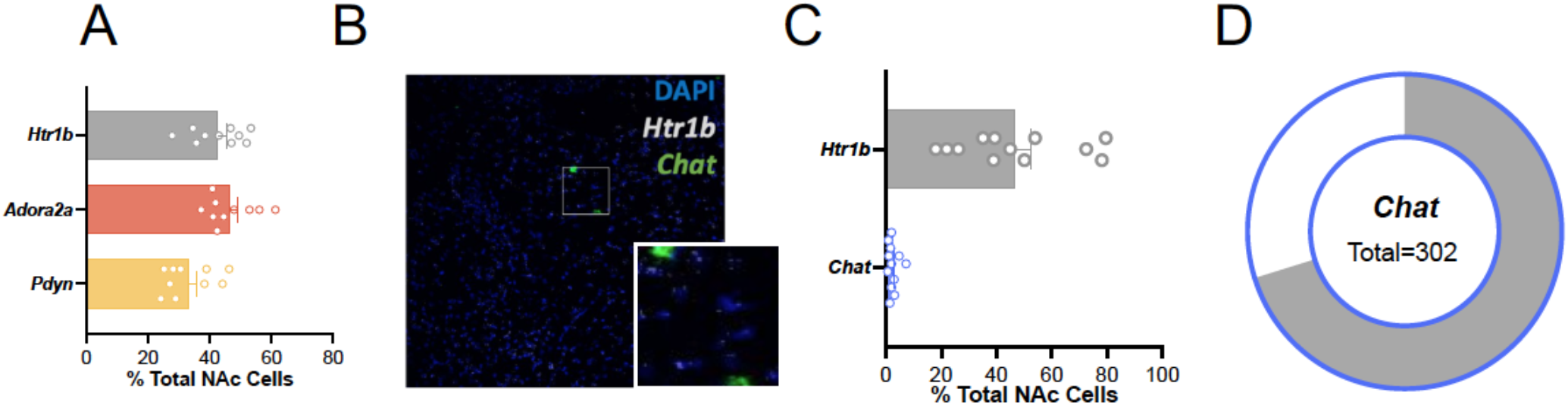
(A) Quantification of percent of total cells in the medial NAc positive for *Htr1b*, *Pdyn*, and *Adora2a* (n=6). (B) Representative image showing *in-situ* hybridization labeling *and Htr1b* and *Chat* transcripts in the medial NAc. Inset: higher magnification of rectangular region. Scale bar=100μm, 25μm. (C) Quantification of percent of total cells in the medial NAc positive for *Htr1b* and *Chat* (n=6). (D) Fraction of *Chat* cells expressing *Htr1b*.

## Notes

### Competing Interest Statement

The authors have declared no competing interest.

